# CRISPRa screen on a genetic risk locus shared by multiple autoimmune diseases identifies a dysfunctional enhancer that affects *IRF8* expression through cooperative lncRNA and DNA methylation machinery

**DOI:** 10.1101/2021.06.11.448156

**Authors:** Tian Zhou, Xinyi Zhu, Zhizhong Ye, Yongfei Wang, Chao Yao, Ning Xu, Mi Zhou, Jianyang Ma, Yuting Qin, Yiwei Shen, Yuanjia Tang, Zhihua Yin, Hong Xu, Yutong Zhang, Xiaoli Zang, Huihua Ding, Wanling Yang, Ya Guo, John B. Harley, Bahram Namjou, Kenneth M. Kaufman, Leah C. Kottyan, Matthew T. Weirauch, Guojun Hou, Nan Shen

## Abstract

Dysregulated transcription factors represent a major class of drug targets that mediate the abnormal expression of many critical genes involved in SLE and other autoimmune diseases. Although strong evidence suggests that natural human genetic variation affects basal and inducible gene expression, it is still a considerable challenge to establish a biological link between GWAS-identified non-coding genetic risk variants and their regulated gene targets. Here, we combine genetic data, epigenomic data, and CRISPR activation (CRISPRa) assays to screen for functional variants regulating *IRF8* expression. Using CRISPR-mediated deletion and 3D chromatin structure analysis, we demonstrate that the locus containing rs2280381 is a cell-type-specific distal enhancer for *IRF8* that spatially interacts with the *IRF8* promoter. Further, rs2280381 mediates *IRF8* expression through enhancer RNA *AC092723.1*, which recruits TET1 to the *IRF8* promoter to modulate *IRF8* expression by affecting methylation levels. The alleles of rs2280381 modulate PU.1 binding and chromatin state to differentially regulate *AC092723.1* and *IRF8* expression. Our work illustrates a strategy to define the functional genetic variants modulating transcription factor gene expression levels and identifies the biologic mechanism by which autoimmune diseases risk genetic variants contribute to the pathogenesis of disease.

## Introduction

Transcription factors (TFs) are specialized proteins that bind to sequence-specific DNA sequences to activate or inhibit gene transcription^1^. TFs contribute to the control the gene-expression pattern of each cell type and the determination of cell fate^2,3^. Emerging evidence reveals that abnormally expressed TFs can contribute to dysregulation of the immune system, and ultimately to the development of autoimmune diseases in mice and humans^3–5^. Studies using TF knockout mice, such as *IRF5*, have directly revealed a critical role for TFs in the pathogenesis and severity of autoimmune diseases^6^. Moreover, inhibition of TFs can effectively intervene disease development, making TFs attractive therapeutic targets in many diseases^7–9^.

The expression levels of many genes differ among individuals, with genetic variants likely making important contributions^10–12^. In particular, genetic variants can alter the expression of genes encoding TFs, thus resulting in alterations to the downstream expression levels of genes controlled by a particular TF^13–15^. For example, *BCL11A* plays a key role in the repression of γ-globin expression and fetal hemoglobin in erythroid cells. Genome-wide association studies (GWAS) have identified genetic variants in the *BCL11A* locus that are associated with fetal hemoglobin expression levels, and targeting *BCL11A* can prevent or ameliorate the complications of sickle cell disease by regulating γ-globin expression levels^15,16^. Dissecting the impact of functional genetic variants on TF expression thus can help to shed light on the mechanisms underlying abnormal expression of transcription factors in disease, especially for diseases with a genetic predisposition.

Autoimmune diseases are a class of complex heterogeneous disease that are broadly caused by an immune response against self^17^. Many autoimmune diseases, such as systemic lupus erythematosus (SLE), have genetic predisposition^18–20^. Abnormal expression of transcription factors leading to the dysregulation of multiple signaling pathways is thought to extensively contribute to autoimmune disease development^21,22^. However, little autoimmune disease risk genetic variants have been directly connected to transcription factor expression levels. This is because genetic variants identified through GWAS are not necessarily causal due to linkage disequilibrium (LD)^18^. Moreover, the majority of variants are located in non-coding genomic regions, and thus are more likely to act in a cell or context-dependent manner^23–26^. In addition, disease risk genes are usually defined based on genomic proximity or expression quantitative trait loci (eQTL) signal, which do not necessarily identify the causal genes.

To fill this gap, we present a strategy for deciphering the mechanism of genetic regulation of transcription factor expression in diseases, using *IRF8* as an example. *IRF8* has been nominated as an important autoimmune disease risk gene by multiple genetic studies^27–34^. Consistent with this notion, *IRF8* function is linked to multiple autoimmune-related phenotypes, such as immune cell development, inflammatory cytokine production and regulation of IFN-stimulated gene (ISG) expression^35–37^. Despite the presence of strong genetic associations in the *IRF8* locus, the functional variants, causal genes, and underlying gene regulatory mechanism involved in autoimmune disease are largely unclear. Here, we integrated genetic data, epigenomic analysis, CRISPR activation (CRISPRa) screens, CRISPR-mediated knockout and 3D chromatin structure analysis to identify the functional variants in the *IRF8* locus. We demonstrate that rs2280381 is likely a causal variant that regulates *IRF8* expression by modulating enhancer RNA expression and cell-type specific enhancer-promoter looping interactions. Furthermore, the enhancer RNA interacts with TET1, which binds to the *IRF8* promoter and modulates its methylation levels to regulate *IRF8* expression. In particular, rs2280381 alleles differentially affect transcription factor occupancy and chromatin state to fine-tune the expression of *IRF8*, thus contributing to disease pathogenesis.

## Results

### CRISPR activation screen identifies the functional autoimmune diseases associated genetic variants in the *IRF8* locus

*IRF8* locus is strongly associated with autoimmune diseases, including Bechet’s disease (BD)^34^, rheumatoid arthritis (RA)^38,39^, systemic sclerosis (SSc)^40,41^, systemic lupus erythematosus (SLE)^28,29,32,42^ and multiple sclerosis (MS)^27,43,44^. GWAS have reported at least 16 genetic variants that are genome-wide significantly associated with autoimmune disease in this locus^27–29,32,34,38–44^. However, the functional variants are currently unknown. To prioritize autoimmune diseases risk variants with potential regulatory function on *IRF8* expression, we developed a strategy to screen the genetic variants with CRISPR activation assays using gRNAs targeting the SNP-containing region. We first collected all autoimmune disease associated genetic variants with genome-wide significance (*P* < 5×10^−8^) published through 2020^27–29,32,34,38–44^ (Fig. 1A and Supplementary Data Set 1). To include all possible disease-associated variants, we further included all SNPs in tight LD (r^2^>0.8) with these tag variants according to the population, identifying 89 SNPs in total (Supplementary Data Set 1). Since GWAS variants are mostly located in non-coding regions of the genome, and variants impacting gene regulation are often located within enhancer regions, we analysed active enhancer signals including H3K27ac modification, DNase I hypersensitive sites (DHSs) and chromatin accessibility (ATAC-seq) of the above SNPs in 4 major human immune cell subsets. SNPs with high H3K27ac, DNase-seq, or ATAC-seq signal in any immune cell subset were considered as candidates (Fig. 1B). This procedure identified 32 candidate genetic variants (Supplementary Data Set 1). Among these SNPs, 18 SNPs are located in monocyte-specific enhancers, with the remaining SNPs mostly occurring in shared enhancer regions of CD4+ T cells, CD8+ T cells, CD19+ B cells and CD14+ Monocytes (Supplementary data set 1 and Fig. 1C). Interestingly, nearly all of these candidate SNP-containing regions are enhancers in CD14+ monocytes (Supplementary Data Set 1 and Fig. 1C). Based on the above observations, we decided to perform our functional screen assays in monocytes.

**Fig. 1.**
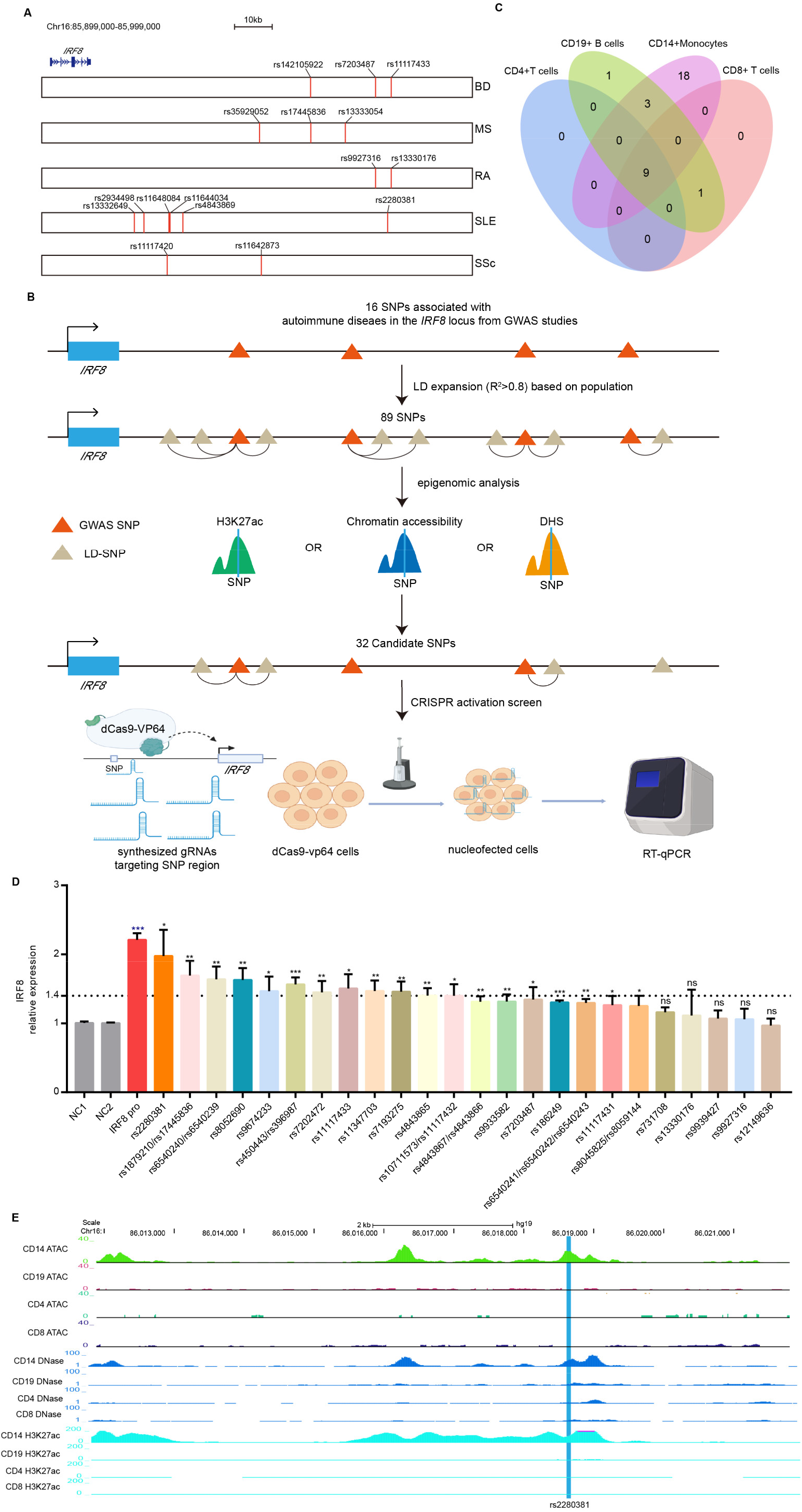
CRISPR activation screen reveals functional genetic variants modulating *IRF8* expression. **(A)** The position of GWAS tag SNPs (shared x-axis indicated above) with respect to the *IRF8* gene for many autoimmune diseases (y-axis). **(B)** Strategy for choosing candidate autoimmune disease-associated SNPs for the CRISPR activation screen. DHS, DNase I hypersensitive site. **(C)** Venn diagram indicating the overlap of SNPs with gene regulatory region signals among different human immune cell subsets. **(D)** RT-qPCR analysis of *IRF8* expression in the CRISPR activation experiment (n = 3, biological replicates). **(E)** Chromatin landscape analysis reveals that rs2280381 is located within a likely cell-type-specific enhancer. Data are represented as mean ± SEM and *P*-values are calculated using an unpaired two tailed Student’s t-test. \**P* < 0.05; \*\**P* < 0.01; \*\*\**P* < 0.001, ns, no significant difference.

To begin functionally identifying the regulatory potential of these SNPs, we first established stable cells expressing dCas9-VP64 in U-937 monocyte cells. Three gRNAs around each SNP were designed and synthesized. The gRNA mixture was then transfected into the cells for 24 hours and *IRF8* mRNA expression levels were measured (Fig. 1B). The results show that 13 variant-harbouring regions could induce greater than 1.4-fold increases in *IRF8* expression levels, with the SLE risk SNP rs2280381-containing region having the strongest regulatory effect among these SNPs (1.97-fold, which is close to the effect of activating the *IRF8* promoter region) (Fig. 1D). Moreover, rs2280381 is located within a monocyte-specific enhancer (Fig. 1E). Based on these results, we focused our study on rs2280381 and SLE.

### The rs2280381-containing enhancer regulates *IRF8* expression in a cell-type-dependent manner through an enhancer-promoter connection

CRISPR/Cas9 mediated deletion is a widely used tool for the study of enhancer function. To directly evaluate the regulatory function of the rs2280381-containing region, we generated cell clones with a roughly 138-bp deletion at the rs2280381 site using the CRISPR/Cas9 technology in U-937 cells (Fig. 2A and 2B). The clones underwent the same procedure, but with the wildtype genotype used as a control. As expected, deletion of the fragment harboring rs2280381 dramatically reduced *IRF8* expression, both at the mRNA and protein level (Fig. 2C and 2D). Further, we also examined the enhancer mark signals in this region by ChIP-qPCR (Fig. 2E) and chromatin accessibility by FAIRE-qPCR in U-937 cells (Fig. 2F), revealing that this chromatin region is open and highly modified with H3K27ac and H3K4me1 marks. These data together indicate that the rs2280381-containing region is a functional enhancer regulating *IRF8* expression in U-937 cells.

**Fig. 2.**
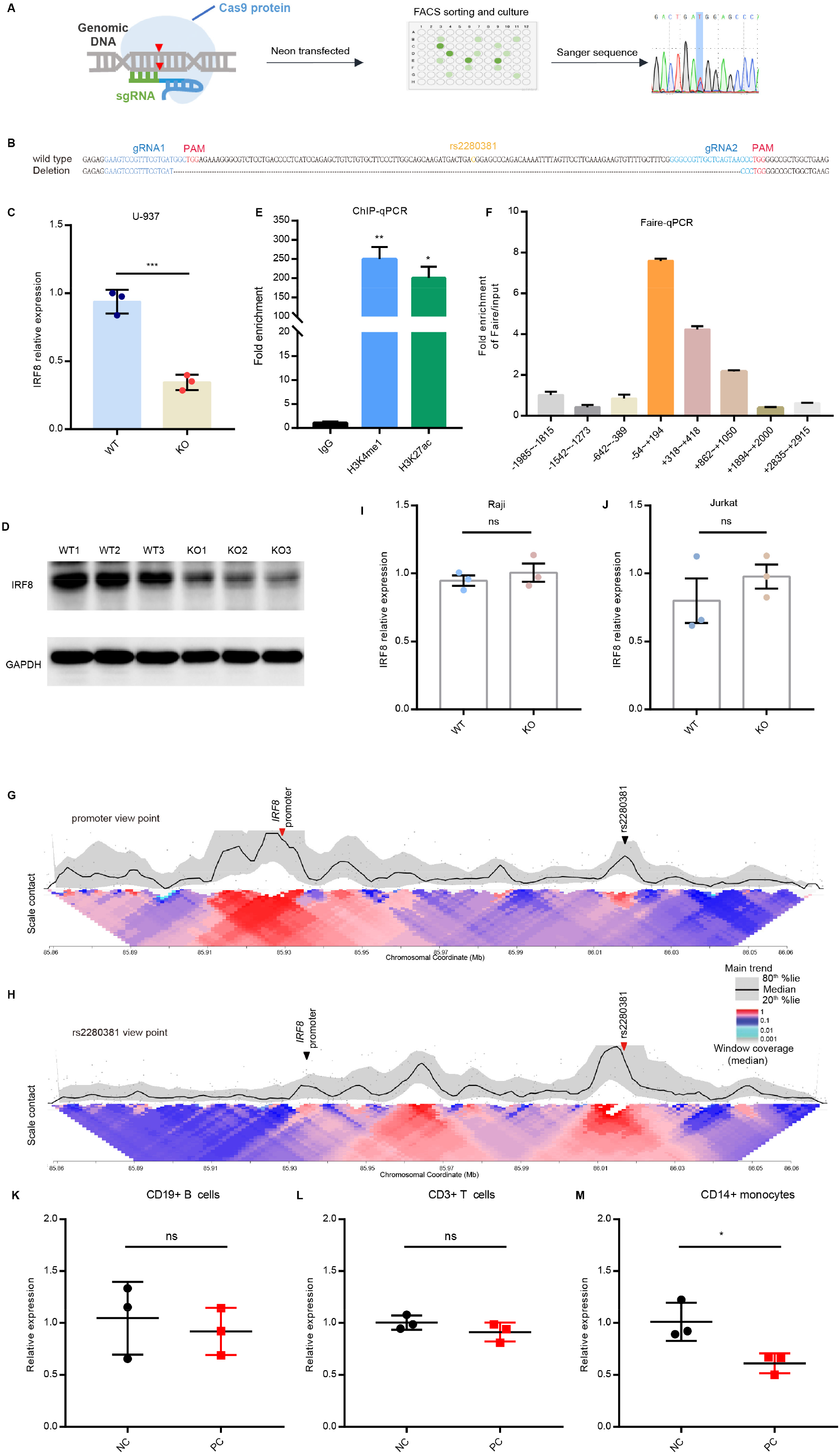
The rs2280381-containing region is a cell-type-dependent enhancer regulating *IRF8* expression. **(A)** Flow chart for generating genomic fragment deletion clones using the CRISR-Cas9 technology. **(B)** The genotype of rs2280381 wildtype clones and deletion clones. **(C)** RT-qPCR analysis of *IRF8* expression in U-937 WT and KO clones (n = 3, biological sample replicates). WT: rs2280381 wildtype, KO: 138 bp fragment harbouring the rs2280381 deletion. **(D)** WB analysis of *IRF8* expression in U-937 WT and deletion clones. WT: rs2280381 wildtype, KO: 138 bp fragment harbouring the rs2280381 deletion. **(E)** Analysis of active enhancer signals (H3K4me1 and H3K27ac) within the rs2280381-containing region in U-937 cells by ChIP-qPCR (n = 3, biological replicates). **(F)** FAIRE-qPCR analysis of chromatin accessibility within the rs2280381-containing region. (n = 3, biological replicates). **(G-H)** 4C-seq analysis of contact profiles of the *IRF8* promoter site (G) and rs2280381 site (H) using a 2 kb window size in the main trend subpanel. Red arrow heads indicate the view point position, and black arrow heads indicate the target position. Gray dots indicate normalized contact intensities. Heat map displays a set of medians of normalized contact intensities calculated at different window sizes. **(I-J)** RT-qPCR analysis of *IRF8* expression in Raji (I) or Jurkat (J) WT and KO clones (n = 3, biological sample replicates). **(K-M)** RT-qPCR analysis of *IRF8* expression in CRISPR/Cas9 RNP edited CD19+ B cells (K), CD3+ T cells (L) and CD14+ monocytes (M) (n = 3, biological samples replicates). Data are represented as mean ± SEM and *P*-values are calculated using an unpaired two tailed Student’s t-test (C, E-F, I-J) and paired two tailed Student’s t-test (K-M). \**P* < 0.05; \*\**P* < 0.01; \*\*\**P* < 0.001.

Distal enhancers usually form enhancer-promoter loops affecting target gene expression. To test whether such a connection exists between the *IRF8* promoter and the rs2280381 enhancer, we conducted circularized chromosome conformation capture sequencing (4C-seq) assays to detect looping interactions to the *IRF8* promoter within this region. These experiments revealed a physical looping interaction between the rs2280381 enhancer and the *IRF8* promoter (Fig. 2G). In addition, this observation was further corroborated based on the rs2280381 view point (Fig. 2H).

Since enhancers are often cell-type-specific, and our data suggest that the rs2280381 enhancer is a monocyte-specific enhancer, we next sought to define in which cell type this region possesses regulatory function. To this end, we first deleted the rs2280381-containing region in Raji (B cell) and Jurkat (T cell) lines, and found that deletion of this region has no effect on *IRF8* expression (Fig.2I and 2J). Next, we isolated CD14+ monocytes, CD3+T cells and CD19+ B cells from human PBMCs and disrupted the rs2280381 region by delivering Cas9 RNP into these cells. After editing, the cells were collected to extract RNA and genomic DNA. The editing efficacy was estimated using ICE (https://ice.synthego.com/#/). Through analysis of the Sanger sequencing results of the target locus and efficiency, up to >30% sample (Fig. S1) was chosen to inspect gene expression. As shown in Fig.2K-M, disruption of the rs2280381-containing region only affects *IRF8* expression in monocytes. This is consistent with the observed epigenetic modifications (Fig.1E) in these immune cells. Collectively, these data suggest that the genomic region harboring rs2280381 is a cell-type-specific enhancer that forms enhancer-promoter interactions to modulate *IRF8* expression.

### LncRNA *AC092723.1* near rs2280381 functions as an enhancer RNA regulating *IRF8* expression

To identify further downstream targets of the rs2280381-containing region and identify other genes in the locus that might be regulated by the region, we performed RNA-seq on three WT and three KO clones. We then performed differential gene expression analysis, identifying 59 and 199 genes significantly downregulated and upregulated (log2 fold-change of ≥ 1.2 and false discovery rate (FDR) cutoff ≤ 0.05) by deletion of this region, respectively (Fig. S2A and Supplementary data set 2). Gene ontology (GO) analysis revealed that differentially expressed genes are highly enriched in expected biological process such as inflammatory response, response to interferon-alpha, LPS or virus, innate immune response, etc. (Fig. S2B and Supplementary data set 2), which is in concordance with the established functions of IRF8^35^.

Interestingly, examination of the expression of genes within the 1-Mb window of rs2280381 revealed that the expression level of a lncRNA, *AC092723.1*, was also down-regulated in rs2280381 KO cell clones (Fig. S2C). This observation was validated by RT-qPCR analysis (Fig. 3A). In addition, use of the CRISPR SAM activation system targeting the rs2280381-containing region by gRNA strongly upregulated both *AC092723.1* and *IRF8* expression (Fig. 3B-C). *AC092723.1* is located downstream of rs2280381, with the distance between rs2280381 and the 3′ end of *AC092723.1* being approximately 300 bp (Fig. 3D). Epigenomic analysis indicates that the genomic region of *AC092723.1* overlaps with a broad monocyte specific likely enhancer with strong H3K4me1, H3K27ac, and DHS signal and high chromatin accessibility (Fig. S2D). Based on these observations, we hypothesized that *AC092723.1* may act as an enhancer RNA (eRNA) mediating the regulation of rs2280381 on IRF8 expression.

**Fig. 3.**
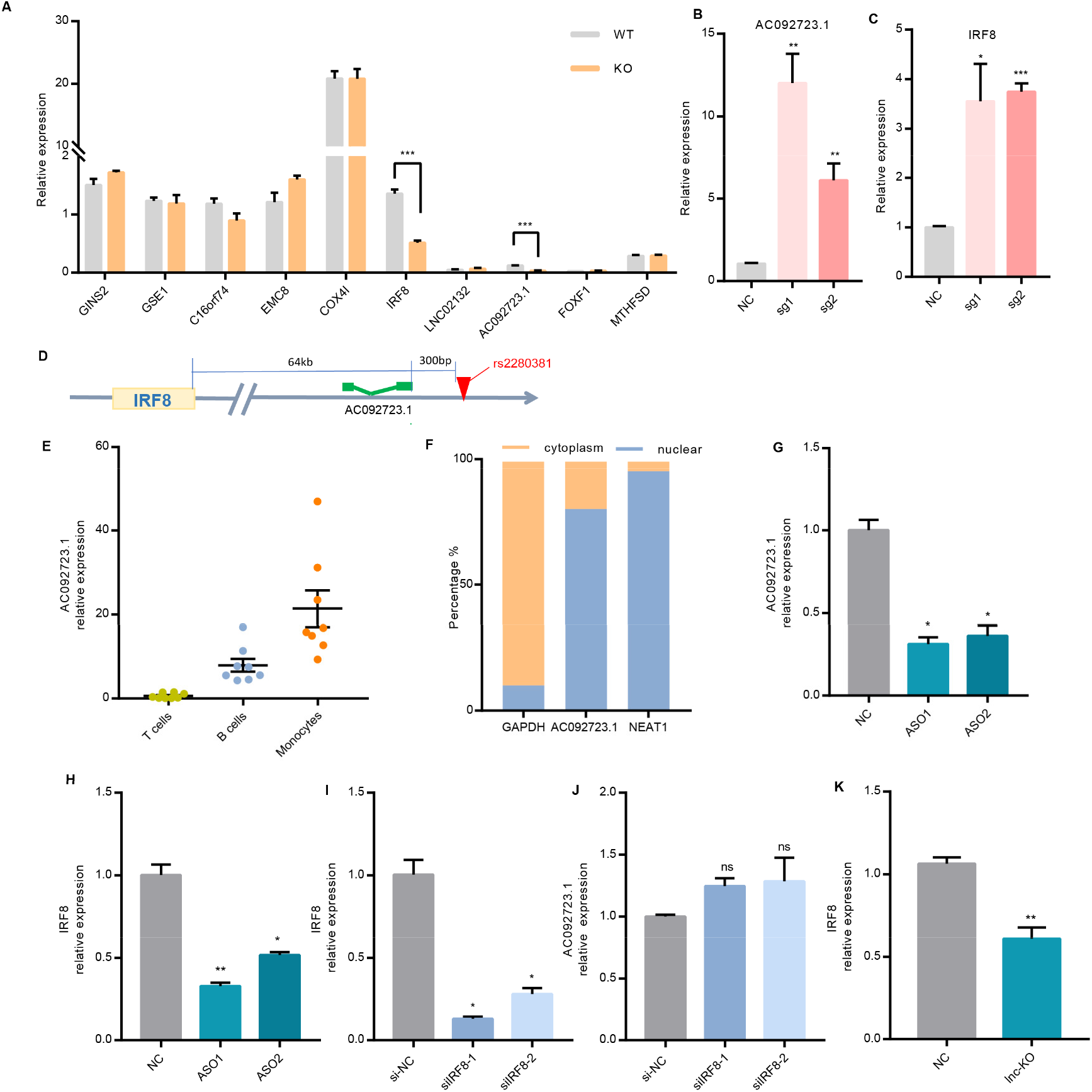
*AC092723.1* mediates the effect of the rs2280381-containing region on *IRF8* expression. **(A)** RT-qPCR analysis of the expression of nearby genes within the 1-Mb window of rs2280381 in U-937 WT clones and KO clones (n = 3, biological sample replicates). **(B-C)** CRISPR SAM assay increases *AC092723.1* and *IRF8* expression by targeting the rs2280381-containing region using specific gRNA in U-937 cells (n = 3, biological replicates). **(D)** The relative locations of rs2280381, *AC092723.1* and *IRF8*. **(E)** *AC092723.1* expression in different human immune cell subsets as measured by RT-qPCR. **(F)** RT-qPCR analysis of *AC092723.1* abundance in U-937 nuclear and cytoplasmic fractions. *GAPDH*, cytoplasmic marker. *NEAT1*, nuclear marker. **(G-H)** RT-qPCR analysis of *AC092723.1* and *IRF8* expression with or without *AC092723.1* knockdown. **(I-J)** RT-qPCR analysis of *AC092723.1* and *IRF8* expression with or without *IRF8* knockdown. **(K)** RT-qPCR analysis of *IRF8* expression after deletion of part region of *AC092723.1* by CRISPR-Cas9. Data are represented as mean ± SEM and *P*-values are calculated using an unpaired two tailed Student’s t-test. \**P* < 0.05; \*\**P* < 0.01; \*\*\**P* < 0.001.

Since lncRNA expression levels are often tissue or cell specific, we first investigated *AC092723.1* abundance in different human immune cell subsets. Consistent with the chromatin landscape in this region, *AC092723.1* is highly expressed in human CD14+ monocytes (Fig. 3E). This was also validated by public RNA-sequencing data in different immune cell subsets (Fig. S2E). Further, we detected its intracellular localization through cell fractionation followed by RT-qPCR in U-937 cells and observed that *AC092723.1* is mainly distributed in the nuclear fraction (Fig. 3F), which is similar to most regulatory lncRNAs. To directly evaluate the regulatory function of *AC092723.1*, we knocked down this lncRNA by ASO and tested *IRF8* expression in U-937 cells. As shown in Fig. 3G-H, knock down of *AC092723.1* significantly reduced *IRF8* expression. In contrast, knockdown of *IRF8* by siRNA did not decrease *AC092723.1* expression (Fig. 3I-J). We further confirmed this result by deleting part of the *AC092723.1* region by CRISPR/Cas9 mediated fragment deletion in U-937 cells (Fig. S2F and Fig. 3K). Collectively, these data provide direct evidence that the rs2280381 enhancer governs eRNA *AC092723.1* expression to modulate *IRF8* expression.

### *AC092723.1* interacts with the TET1 protein and binds to the *IRF8* promoter, regulating *IRF8* expression by influencing methylation levels

To explore the mechanism by which *AC092723.1 cis*-regulates *IRF8* expression, we first carried out a chromatin isolation by RNA purification (ChIRP) assay^45^ to evaluate the interaction between the lncRNA and the *IRF8* promoter. We designed biotinylated antisense oligonucleotides tilling the whole lncRNA sequence and incubated these probes with chromatin fractions from U-937 cells. The core binding sequence within the *IRF8* promoter was assessed by RT-qPCR (Fig. 4A). Five pairs of PCR primers were designed spanning from −1000 to +153 relative to the *IRF8* transcription start site, which include all of the high chromatin accessibility regions (Fig. S3). Compared to the control GAPDH group, lncRNA probes strongly and specifically enriched *AC092723.1* mRNA compared to *GAPDH* mRNA (Fig. 4B-C). More importantly, analysis of the DNA sequences pulled down by *AC092723.1* probes revealed significant enrichment at the −473/-395 *IRF8* promoter sequence (Fig. 4D and Fig. S3). In addition, we also found that *AC092723.1* could interact with the rs2280381-containing region (Fig. 4E), which suggests that *AC092723.1* may contribute to loop formation between the *IRF8* promoter and the rs2280381 enhancer (Fig.4F).

**Fig. 4.**
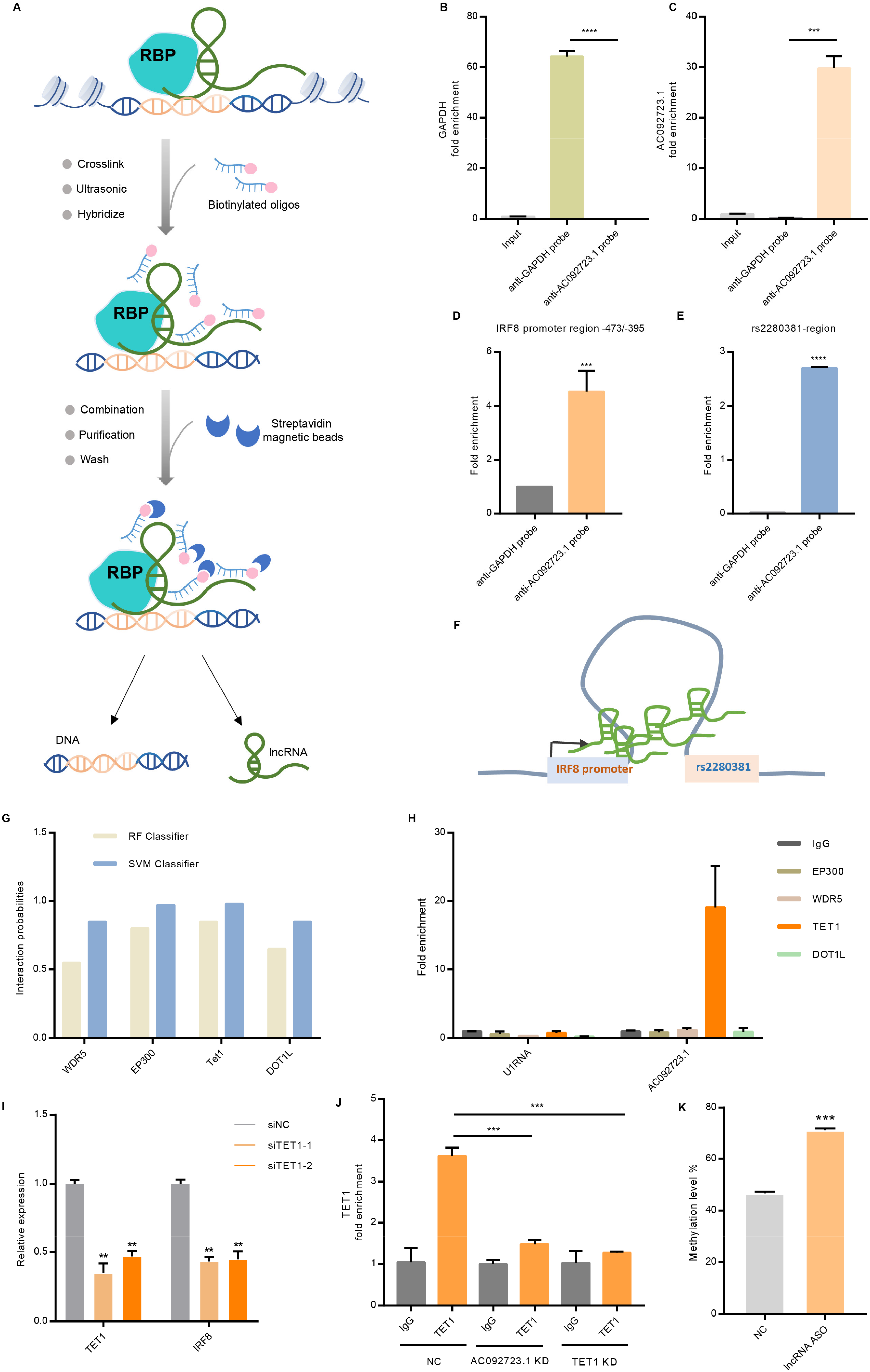
*AC092723.1* binds to the *IRF8* promoter, where it recruits TET1 and affects promoter methylation levels. **(A)** Flow scheme for the ChIRP assay detecting the interaction between the *IRF8* promoter region and *AC092723.1*, RBP: RNA-binding protein. **(B-C)** *GAPDH* mRNA and *AC092723.1* RNA are specifically enriched with anti-GAPDH probes and anti-AC092723.1 probes in ChIRP assays, respectively (n = 3, biological replicates). **(D-E)** *AC092723.1* interacts with the *IRF8* promoter region **(D)** and rs2280381 containing region (E) (n = 3, biological replicates). **(F)** Model for *AC092723.1* contribution to loop formation between the *IRF8* promoter region and the rs2280381 region. **(G)** Bioinformatic analysis of *AC092723.1* interacting chromatin modifiers using online tool RPISeq based on random forest (RF) or support vector machine (SVM) models. Interaction probabilities generated by RPISeq range from 0 to 1, predictions with probabilities > 0.5 indicating that the corresponding RNA and protein are likely to interact. **(H)** RIP-qPCR analysis of the interaction between *AC092723.1* and predicted binding chromatin modifiers (n = 3, biological replicates). **(I)** RT-qPCR analysis of *IRF8* expression in U-937 cells after knockdown of *TET1* by siRNA (n = 3, biological replicates). **(J)** ChIP-qPCR analysis of the binding efficiency of TET1 to the *IRF8* promoter with or without *AC092723.1* knockdown (n = 3, biological replicates). **(K)** Methylation levels of the *IRF8* promoter region in U-937 cells with or without *AC092723.1* knockdown (n = 3, biological replicates). Data are represented as mean ± SEM and *P*-values are calculated using an unpaired two tailed Student’s t-test. \**P* < 0.05; \*\**P* < 0.01; \*\*\**P* < 0.001.

Recent studies have demonstrated that many lncRNAs can function as scaffolds for chromatin-modifying enzymes that regulate chromatin epigenetic modifications to enhance or suppress gene expression^46,47^. To test whether *AC092723.1* can interact with epigenetic modifying enzymes, we first used a bioinformatics algorithm (http://pridb.gdcb.iastate.edu/RPISeq/references.php)^48,49^ to predict possible binding epigenetic modifying enzymes. Since *AC092723.1* is a positive regulator of *IRF8* expression, we focused our candidates on chromatin modifiers with the potential function of activating gene expression: WDR5, EP300, TET1 and DOT1L (Fig. 4G). To test these candidate modifiers, we performed RNA binding protein immunoprecipitation assays (RIP) with antibodies specific to each of the above chromatin modifiers. As shown in Fig. 4H, only the anti-TET1 antibody enriched for high abundance of *AC092723.1* relative to the IgG control.

Next, we investigated whether TET1 plays a functional role with *AC092723.1* in regulating *IRF8* expression. We used siRNA knockdown of *TET1* in U-937 cells and performed RT-qPCR analysis to evaluate the knockdown efficiency and *IRF8* expression levels. The results show that silencing of *TET1* significantly down-regulated *IRF8* expression (Fig. 4I). Further, to elucidate how the *AC092723.1*-TET1 complex modulates *IRF8* expression, we detected the enrichment of TET1 in the *IRF8* promoter region by performing chromatin immunoprecipitation (ChIP) assays in U-937 cells with or without lncRNA knockdown. The results indicate that TET1 can directly bind to the *IRF8* promoter region, and this binding activity is impaired in *AC092723.1* KD cells (Fig.4J), implying that *AC092723.1* functions as a scaffold recruiting TET1 to the *IRF8* promoter region.

TET1 is an important chromatin-modifying enzyme that causes DNA demethylation, thus activating gene expression^50^. To test whether TET1 acts in this fashion to control *IRF8* expression, we examined the methylation level of the *IRF8* promoter region after silencing *AC092723.1* expression. As expected, the methylation level in the promoter region significantly increased upon *AC092723.1* KD (Fig. 4K). Taken together, these data suggest that *AC092723.1* interacts with TET1 to limit the methylation levels in the *IRF8* promoter region, leading to the activation of *IRF8* transcription.

### rs2280381 alleles differentially regulate *AC092723.1* and *IRF8* expression by modulating PU.1 binding and the chromatin state

eQTL data indicate an association between rs2280381 alleles and *IRF8* and *AC092723.1* expression levels (Fig. S4A-B), but there is no direct evidence illustrating that the rs2280381 alleles differentially regulate *IRF8* or *AC092723.1* expression. To investigate whether rs2280381 alleles directly regulate *AC092723.1* and *IRF8* expression, we adopted the prime editing technology^51^ to generate isogenic cell lines carrying rs2280381 homozygous major (T/T), homozygous minor (C/C), or heterozygous (T/C) alleles (Fig. 5A and Fig. S4C). For each genotype, we selected six clones to measure the effect of rs2280381 on *IRF8* and *AC092723.1* expression. In agreement with the eQTL data, RT-qPCR data showed that the SLE risk allele (T) results in lower expression of *AC092723.1* and *IRF8* compared to the non-risk allele (C) (Fig. 5B-C), consistent with the down-regulated expression of *IRF8* and *AC092723.1* in SLE patients (Fig. S4D-E).

**Fig. 5.**
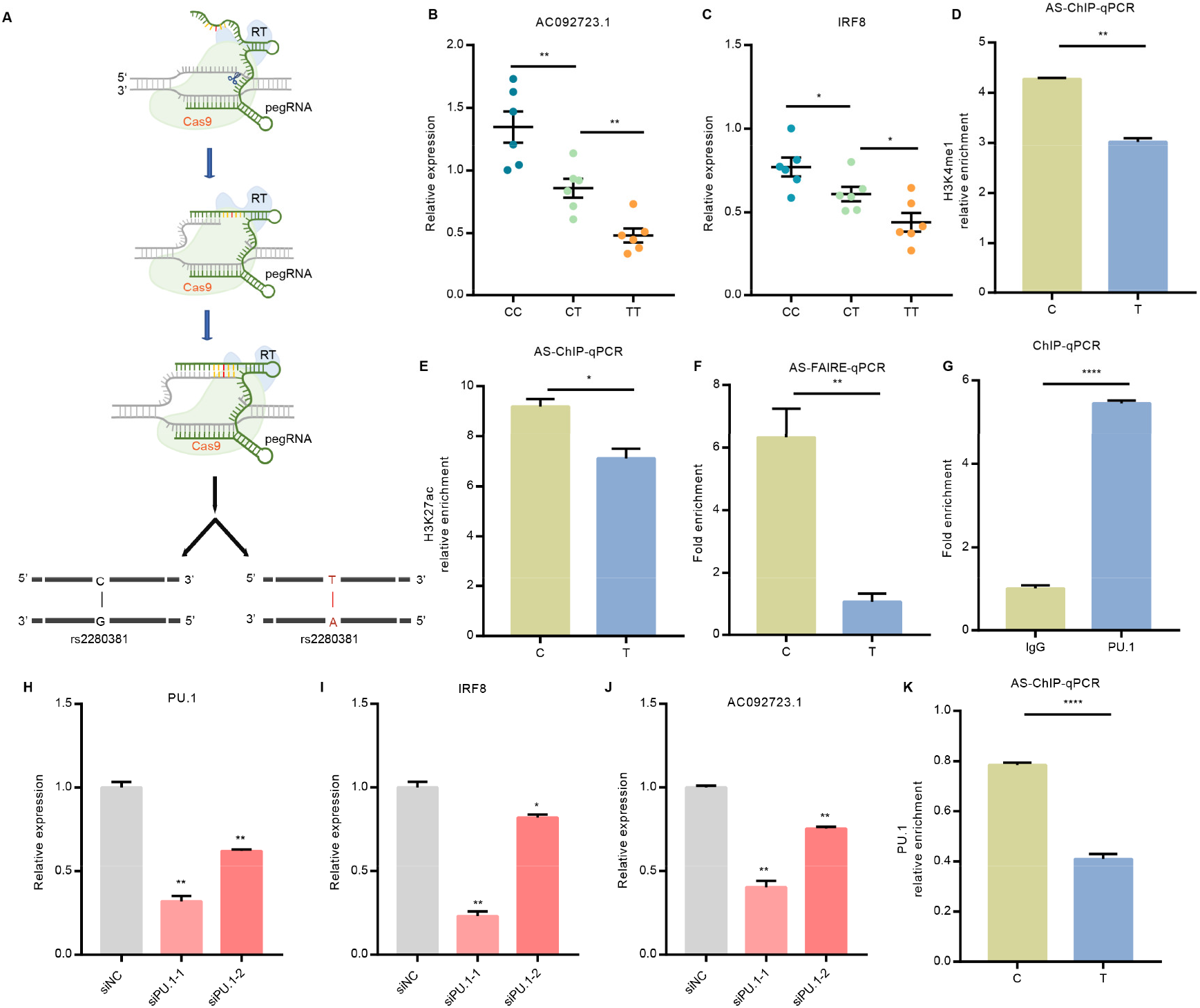
rs2280381 alleles affect H3K4me1, H3K27ac and PU.1 binding to fine tune the expression of *AC092723.1* and *IRF8*. **(A)** Work flow for the generation of isogenic cell clones with the Prime editing technology. **(B-C)** The rs2280381 C allele leads to higher expression of *AC092723.1* and *IRF8* compared to the T allele. (n = 6, biological samples replicates). **(D-E)** H3K4me1 and H3K27ac histone marks are stronger for the rs2280381 non-risk C allele, as determined by AS-ChIP-qPCR in the rs2280381 heterozygous U-937 cell clone. (n = 3, biological replicates). **(F)** The genomic region harboring the non-risk allele (C) exhibits increased chromatin accessibility compared to the risk allele (T), as determined by AS-FAIRE-qPCR in the rs2280381 heterozygous U-937 cell clone. (n = 3, biological replicates). **(G)** Relative enrichment of PU.1 binding to the rs2280381-containing region, as measured by ChIP-qPCR in U-937 cells (n = 3, biological replicates). **(H-J)** Relative expression of *PU.1* (H), *IRF8* (I) and *AC092723.1* (J) after PU.1 siRNA-mediated knockdown, as measured by RT-qPCR in U-937 cells (n = 3, biological replicates). **(K)** PU.1 binds more strongly to the rs2280381 C non-risk allele, as determined by ChIP followed by AS-qPCR in the rs2280381 heterozygous U-937 cell clone (n = 3, biological replicates). Data are represented as mean ± SEM and *P*-values are calculated using an unpaired two tailed Student’s t-test. \**P* < 0.05; \*\**P* < 0.01; \*\*\**P* < 0.001.

After demonstrating the allele-specific regulation capacity of rs2280381, we next sought to explore the underlying mechanisms. Genetic variant changes are often associated with differential enhancer activity, which has been considered as an important mechanism for SNP allele-specific regulation of gene expression. To test whether the different rs2280381 alleles can alter chromatin state, we first carried out H3K4me1 and H3K27ac ChIP experiments followed by allele-specific qPCR in a cell line that is heterozygous for rs2280381. The resulting data reveal that the C allele of rs2280381 contains stronger H3K4me1 and H3K27ac histone mark signals than the T allele (Fig. 5D-E). We also detected the chromatin accessibility of the rs2280381 alleles by FAIRE allele-specific qPCR. In agreement with the allelic bias in histone modifications, enhancers harboring the C allele exhibit more FAIRE signal than enhancers harboring the T allele (Fig. 5F), indicating that the C allele has higher chromatin accessibility than the T allele.

Differential transcription factor binding is another feature of SNP-dependent cis-regulatory effects on gene expression. To determine differential binding of transcription factors to the rs2280381 sequence, we carried out a DNA-affinity precipitation assay (DAPA) followed by mass spectrometry (MS) experiment, revealing 100 candidate proteins. Most proteins identified by DAPA-MS are histone proteins or chromatin structure maintenance proteins (Supplementary Data Set 3). Given that the rs2280381-containing region is a monocyte-specific enhancer, we focused on monocyte-specific transcription factors, identifying PU.1, an important monocyte lineage-determining transcription factor^52^, as our top candidate. ChIP-qPCR experiments were performed, with the results verifying PU.1 binding of the rs2280381-containing region (Fig. 5G). Moreover, siRNA knockdown of PU.1 expression strongly reduced *AC092723.1* and *IRF8* expression (Fig. 5H-J). In addition, we also detected allelic binding of PU.1 to rs2280381, with AS-ChIP-qPCR data indicating that the C allele has stronger PU.1 binding than the T allele (Fig. 5K). Taken together, the enhancer with the rs2280381 C allele has stronger signals for PU.1, H3K4me1, and H3K27ac, and exhibits stronger chromatin accessibility relative to the T allele, enhancing the expression of *AC092723.1* and *IRF8*.

## Discussion

Transcription factors play a critical role in autoimmune disease development, and disease-associated genetic variants are useful for revealing critical mechanisms involved in disease pathogenesis^16,53^. However, only a small number of functional genetic variants have been identified that alter transcription factor gene expression levels. To fill this gap, we designed a general strategy for defining the functional variants regulating *IRF8* expression. Application of this strategy identified rs2280381 as a causal variant in the *IRF8* locus. We also elucidate the specific biological mechanism underlying rs2280381 mediation of SLE risk: alteration of PU.1 binding, histone marks, chromatin accessibility, and lncRNA expression, leading to differential *IRF8* promoter methylation levels and altered *IRF8* expression (Fig. 6).

**Fig. 6.**
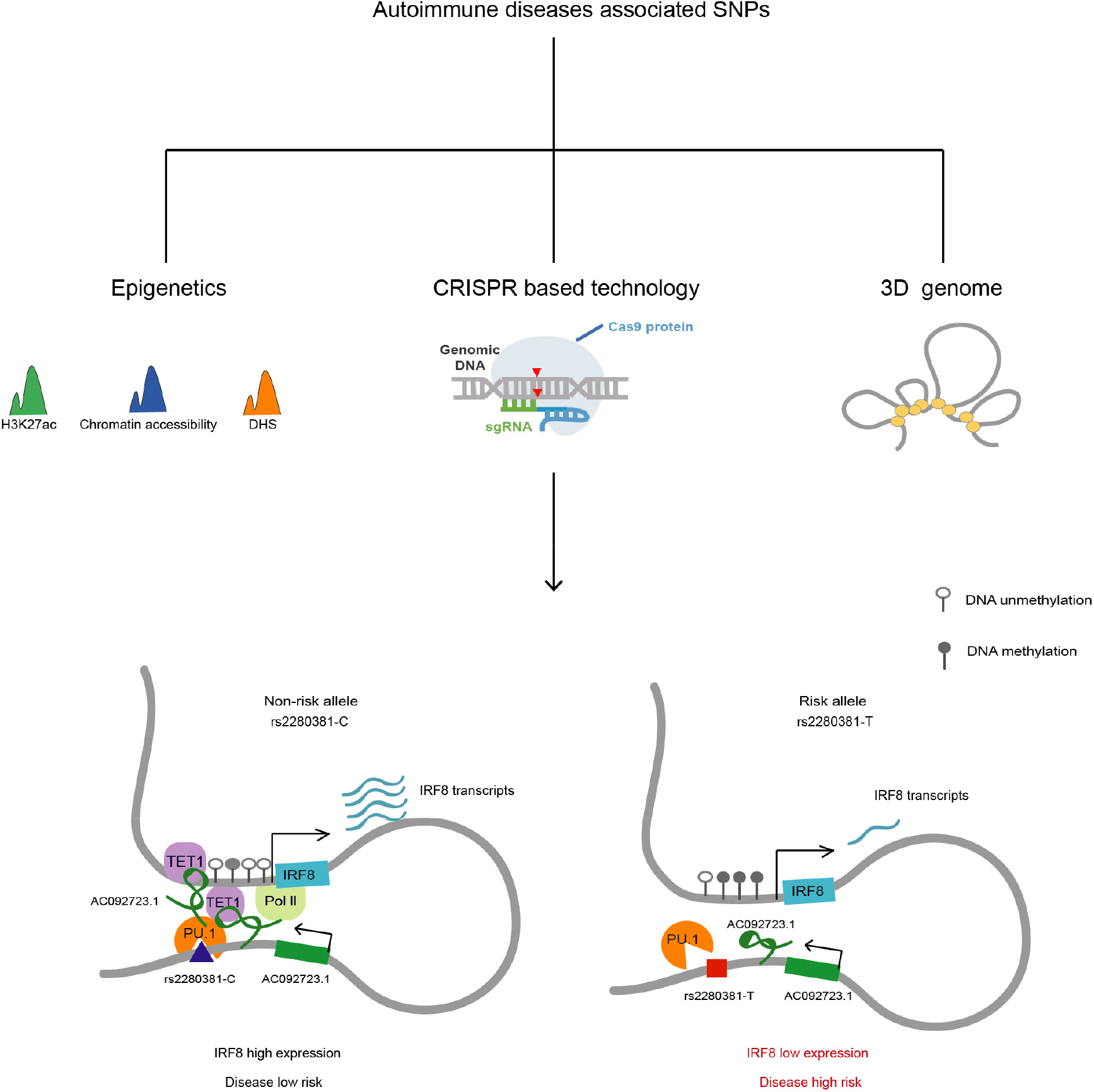
Model for rs2280381 regulated *IRF8* expression mediating disease risk. The rs2280381-containing region forms gene-loop with the *IRF8* promoter region. The rs2280381 T risk allele has lower PU.1-binding affinity than the non-risk C allele, resulting in the reduction of *AC092723.1* expression, upregulation of methylation levels of *IRF8* promoter and decreased *IRF8* expression contributing to SLE risk.

Progress towards discriminating the functional genetic variants regulating transcription factors is continuously challenged by the existence of linkage disequilibrium, the specific cell type(s) where the variant functions, and complications inherent to deciphering gene regulatory mechanisms^18^. Through integration of genetic data with epigenomic analysis, we designed an approach that first ranks all autoimmune disease-associated SNPs in the *IRF8* locus based on the presence of active enhancer histone marks, and then identify candidate SNPs with potential regulatory function. Using CRISPR activation assays, we systematically screened these genetic variants based on their ability to modulate *IRF8* expression, effectively identifying the functional regulatory elements harboring disease-associated SNPs. This strategy provides a blueprint for identifying the functional SNPs regulating the expression of genes encoding transcription factors or other molecules.

Enhancers have been considered effective therapeutic targets for disease intervention because targeting enhancers might aid in precise treatment, due to the cell-type specific nature of enhancers^15,54–56^. For instance, editing the erythroid-specific enhancer of *BCL11A* by CRISPR-Cas9 restores γ-globin synthesis for treating sickle cell disease^15^. Uncovering disease-critical enhancers would thus provide valuable therapeutic targets for disease treatment. In this study, using CRISPR-Cas9 mediated deletion, we edited the rs2280381-containing region in different cell lines and different immune cell subsets and found that the rs2280381-containing region acts as a distal and cell-type-specific enhancer to modulate *IRF8* expression, which suggests that the rs2280381 enhancer has the potential to be a therapeutic target for SLE treatment in the future. In this manner, deciphering the functional genetic variants associated with autoimmune disease will aid in the development of novel treatment methods.

Gene expression is controlled by a series of regulatory elements, including distal enhancers and the proximal promoter^57,58^. Distal enhancers spatially interact with promoter regions to regulate target gene expression^59^. We performed 4C-seq assays to verify promoter-enhancer loops between the *IRF8* promoter and the rs2280381-containing region, which further supports the regulatory function of the rs2280381-containing region. Interestingly, we also observed connections between the *IRF8* promoter site and various other genomic regions (Supplementary Data Set 4), some of which also contain autoimmune disease-associated genetic variants. Although regulatory function for these variants has been validated in the CRISPRa screen assays, the function of these regions remains unknown. Dissecting the function of these regulatory elements will likely aid in our understanding of the complete picture of *IRF8* transcriptional regulation.

A major challenge inherent to the study of non-coding genetic variants is the verification of the functional consequences of the different alleles on gene expression. In this study, we generated cell clones harboring the two rs2280381 alleles by prime editing and demonstrated allele-specific regulation of rs2280381 on *IRF8* expression. However, most individual SNPs only have a small effect on gene expression or disease-associated phenotypes. Some studies have discovered that genetic variants within multiple enhancers of a gene could synergistically regulate gene expression, thus amplifying these individually small effects^60^. In our study, in addition to the rs2280381-containing region, we found several other genetic variant-containing regions, such as rs8052690, that also could increase *IRF8* expression in the CRISPR activation screen assay. These data suggest that the combination of functionally independent genetic variants may be an important risk factor for disease. To fully uncover the mechanism of genetic-mediated disease risk, the synergistic effect of multiple genetic variants should be emphasized in future studies.

Allele-dependent transcription factor binding is a major contributor to allelic gene expression differences. Using DAPA-MS data and ChIP-qPCR, we found that PU.1 is the key transcription factor binding to the rs2280381 site, and that PU.1 binds differentially to the rs2280381 non-risk allele and risk alleles, which likely leads to the differential regulatory function of the risk and non-risk alleles. The accessibility of transcription factor binding sites is significantly heterogeneous in human immune cells, monocytes exhibited high activity of PU.1^61^, and PU.1 is a key lineage-determining TF for priming monocyte-specific enhancers^52^, the binding of PU.1 to the rs2280381 locus may contribute to its function as a cell-type-specific enhancer. We also observed differential chromatin states for the risk and non-risk alleles, which was reflected by the high H3K27ac enrichment, H3K4me1 enrichment and chromatin accessibility observed for the non-risk C allele compared to the T risk allele. Collectively, these observations elucidate the mechanism underlying rs2280381 risk allele-mediated disease risk. However, whether other proteins involved in this allele specific regulation and the mechanism forming cell-type-specific enhancer still deserve to be studied in more depth.

Intriguingly, our results indicate that the enhancer RNA *AC092723.1* is involved in the regulatory mechanisms underlying the differential effect of the rs2280381 alleles on *IRF8* expression. We show that the rs2280381 alleles are associated with *AC092723.1* expression differences, and *AC092723.1* can directly regulate *IRF8* expression levels. LncRNA can participate in chromatin remodeling complexes that modify the chromatin to enhance or suppress gene expression^46,47^. Using a combination of bioinformatics-based prediction and RIP-qPCR assays, we found that TET1 is a binding partner of *AC092723.1*. TET1 is a key chromatin modifier that modulates gene expression by influencing DNA methylation levels^50^. Our ChIRP and ChIP assays demonstrated that *AC092723.1* recruitment of TET1 results in TET1 binding to the *IRF8* promoter, reducing DNA methylation levels, and thus modulating *IRF8* expression. DNA methylation changes contribute to SLE pathogenesis, but the factors regulating methylation in SLE are largely unknown. Our study links an SLE-associated genetic variant to DNA methylation and ultimately SLE etiology, adding another layer of regulation for genetic variant-based modulation of gene expression involved in the disease.

In conclusion, our study provides a blueprint for the establishment of a link between disease risk genetic risk variants and transcription factor gene expression levels, and applies this approach to decipher an important mechanism underlying SLE risk SNP-mediated disease pathogenesis. Our work also provides key insights that form a strong foundation for the development of disease therapies based on enhancer modulation.

## Methods

### Cell culture

All cell lines were purchased from the Chinese academy of science cell bank (Shanghai, China). U-937, Raji and Jurkat were cultured in 10% (v/v) fetal bovine serum (FBS) and 90% RPMI-1640 medium. HEK-293T was cultured with 10% (v/v) FBS and 90% Dulbecco’s Modified Eagle Medium (DMEM). Cells were maintained at 37 °C and 5% CO_2_ constant temperature incubator. These cell lines were free of mycoplasma during our study.

### RNA extraction and RT–qPCR

Total RNA was extracted from cells using TRIzol Reagent (Invitrogen). Samples of 500 ng of RNA were reverse transcribed using PrimeScript™ RT Reagent Kit (Perfect Real Time) (TAKARA, RR037A). qPCR was performed using TB Green Premix Ex Taq reagent (TAKARA, RR420A). GAPDH expression was determined as an internal control and fold change in expression level was calculated using the ΔΔCt method.

### Western Blotting

Protein lysates were separated on 10% SDS/PAGE gels, transferred to PVDF membranes and probed with antibodies directed against IRF8 (Cell Signalling Technology, 5628S, 1:1000 dilution), GAPDH (Abcam, AC035, 1:5000 dilution). GAPDH was used as a loading control.

### ASOs and siRNAs transfection

Antisense oligonucleotides (ASOs) and siRNAs were synthesized by Sangon Biotech (Shanghai, China). Before transfection, 2×10^5^ cells were seeded into a 24-well plate and incubated at 37 °C and 5% CO_2_ overnight. Next, 200 nM of ASO or siRNA were transfected into the cells using TransIntroTM EL Transfection Reagent (Transgene, FT201-01) and cells were collected to extract RNA.

### RNA Immunoprecipitation (RIP)-qPCR

RIP assays were performed using the EZ-Magna RIP Kit (Millipore). 1×10^7^ cells were lysed with RIP lysis buffer. Cell extracts were coimmunoprecipitated with Anti-TET1 antibody (Abcam, ab191698), Anti-KAT3B/p300 antibody (Abcam, ab10485), Anti-WDR5 antibody (Abcam, ab56919), and Anti-DOT1L Antibody (Thermo, A300-953A). The recovered RNA was subjected to RT-qPCR analysis and U1 was used as a nonspecific control target.

### Candidate SNP picking

LD expansion was done by the online tool (https://pubs.broadinstitute.org/mammals/haploreg/haploreg.php) to include all SNPs in strong LD (r^>0.8) with the reported tag SNPs at the *IRF8* locus. The chromatin landscape of SNP-located regions was analyzed using the public resource provided by the NIH Roadmap Epigenomics Mapping Consortium (http://www.roadmapepigenomics.org/). And SNP-located regions with any signal of ATAC-seq peaks, H3K27Ac peaks or DNase peaks in four major human immune cell subsets were selected as candidate SNPs to undergo CRISPRa screening assays.

### Cell fractionation

This assay was performed using the Nuclear and Cytoplasmic Extraction Kit (CWBIO, Shanghai). In brief, 1 × 10^7^ cells were harvested and resuspended in 1 mL of Nc-buffer A and 55 μL of Nc-buffer B and incubated on ice for 10 min. Cells were then centrifuged for 15 min at 12,000 × g and the supernatant was collected as the cytoplasmic fraction. The pellets were resuspended for 40 min on ice in 500 μL of Nc-buffer C supplemented with RNase inhibitors, centrifuged for 15 min at 12,000 × g and the supernatant was collected as the nuclear fraction. All fractions were resuspended in TRIzol to extract RNA.

### Chromatin Immunoprecipitation (ChIP)-qPCR

This assay was performed using the SimpleChIP® Plus Enzymatic Chromatin IP Kit (Cell Signaling Technology). Briefly, 5×10^6^ cells were first cross-linked by 1% formaldehyde solution and then quenched by 125 mM glycine solution. After that, cells were washed by cold PBS for twice. Cell pellets were resuspended with 1 mL cold 1× Buffer A, incubated on ice for 10 min and centrifuged to remove supernatant. Then the pellets were resuspended in 1 mL cold 1×Buffer B, centrifuged to remove supernatant and resuspended in 100 μL 1×Buffer B. 0.5 μL of Micrococcal Nuclease was added and incubated at 37 °C for 20 min to digest DNA into 150-900 bp length. 10 μL of 0.5 M EDTA was added to stop the digestion, then samples were centrifuged to discard the supernatant. Finally, pellets were resuspended with 100 μL of 1×ChIP Buffer, incubated on ice for 10 min and subsequently sonicated at 4 °C with a Bioruptor sonicator (Diagenode) at high power for 5 cycles with 30 s ON and 30 s OFF. This was centrifuged and supernatant was collected into a new tube and incubated with anti-H3K27 antibody (ab177178, Abcam, 2 μg for 25 μg of chromatin) or anti-H3K4me1 antibody (ab8895, Abcam, 2 μg for 25 μg of chromatin) or anti-PU.1 antibody (2266S, Cell Signaling Technology, 1:50) overnight at 4 °C on rotation. ChIP-grade protein A+G magnetic beads (Millipore, 16-663) were added and the enriched chromatin was eluted with 150 uL ChIP Elution Buffer. DNA fragments were purified with spin columns and enrichment was detected by qPCR.

### Formaldehyde-Assisted Isolation of Regulatory Elements (FAIRE)-qPCR

1×10^7^ cells were cross-linked by 1% formaldehyde solution and then quenched by 125 mM glycine solution. Cells were then sonicated, equal volume of phenol/chloroform/isoamyl alcohol was added into the chromatin lysate and centrifuged to isolate the aqueous. The aqueous was further purified by adding chloroform/isoamyl alcohol. Then DNA was precipitated, washed and reverse crosslinked to prepare the FAIRE DNA. FAIRE DNA samples were analyzed by quantitative RT-PCR with specific primers targeting DNA sequences at different distances to rs2280381 site. Values were normalized to input DNA and compared to a region just outside the putative regulatory region.

### Allele-specific qPCR

We designed AS-qPCR primers to specifically amplify the rs2280381 region with a T or C allele in the ChIP or FAIRE DNA samples. AS-qPCR was performed according to normal qPCR procedures.

### RNA library preparation, sequencing and gene expression analysis

Total RNA was extracted using TRIzol Reagent. rRNA was depleted from total RNA using Ribo-Zero™ rRNA removal Kit and library was made using Illumina NEBNext® Ultra™ Directional RNA Library Prep Kit (E7420L, NEB). The libraries were loaded on an Illumina HiSeq X ten instrument (Illumina). Sequencing was carried out using a 2×150 paired-end configuration. Computational analysis of paired-end reads was conducted using cutadapt (v1.15), Samtools (v0.1.19), Hisat2 (v2.1.0), and HT-seq (v0.11.2) software. Statistical normalization and differential analysis were performed in R using the DESeq2 (v1.24.0) package. The threshold to define up or down regulation was log2 fold-change > 1.2 and P value < 0.05. Visualization was also conducted in R (v3.3.3).

### Chromatin isolation by RNA purification (ChIRP)-qPCR

Probes were designed using an online tool (http://singlemoleculefish.com). Oligonucleotides were synthesized and biotinylated at the 3’ end. 2×10^7^ cells were first cross-linked in 1% glutaraldehyde solution at room temperature, then quenched by 1/10 volume of 1.25 M glycine. Pellets were washed by cold 1×PBS, resuspended in ChIRP lysis buffer and sonicated into 100-500 bp length. 20 μL of lysate was removed to prepare input RNA and DNA sample, then 2 mL hybridization buffer and total 100 pmol probes were added to the remaining lysate and incubated at 37 °C with gentle shaking for 4 hours. 100 μL Dynabeads™ MyOne™ Streptavidin C1(Invitrogen, 65001) were added into each tube, incubated at 37 °C with gentle shaking for 0.5 hour to isolate the chromatin. Chromatin samples for isolating RNA were treated with proteinase K, boiled at 95 °C, then quickly chilled on ice and RNA was extracted using 1 mL TRIzol. RNA samples were then reverse transcribed into cDNA. *AC092723.1* and *GAPDH* enrichment was detected by RT-qPCR, respectively. Chromatin samples for isolating DNA were treated with RNaseA and proteinase K, and purified with phenol/chloroform/isoamyl. DNA samples were directly utilized as a template to detect the enriched region.

### Genome editing in cell lines

For Prime editing, pegRNA was designed using the online CRISPR tool (http://pegfinder.sidichenlab.org/). For constructing nicking gRNA expression vector, pKLV-U6gRNA(BbsI)-PGKpuro2ABFP(Addgene, 50946) was linearized by BbsI (NEB, R3059L) and then gel-purified. Guide RNA oligos were synthesized in Tsingke (Shanghai, China), annealed and subcloned into the linearized 50946 plasmid and were transformed into chemically competent Escherichia coli (Stbl3, Transgen Biotech) to extract plasmid DNA. For constructing pegRNA expression vector, 50946 plasmid was cut by BbsI (NEB, R3059L) and BamHI (NEB, R0136S) and then gel-purified. Guide RNA oligos, gRNA scaffold oligoes, RT temple and prime binding sequence oligoes were annealed and subcloned into the BbsI and BamHI cut plasmid. For editing, 2×10^6^ U-937 cells were prepared and washed by PBS for electro-transfection, 10 μg pCMV-PE2-P2A-GFP (Addgene, 132776) plasmid, 10 μg plasmid expressing pegRNA and 5 μg plasmid expressing nicking gRNA were added to cells and resuspend with 100 μL buffer R, then cells were transfected with the condition of 1400 v, 10 ms, 3 pulses using Neon system. Cells were immediately plated to 6-well plate and cultured for 72 hours. Single cell with strong GFP and BFP signals was sorted into the 96-well plate containing 200 μL culture medium in each well by FACS. After 14 days culture, clones were transferred to a 24-well plate and genotype was identified by sanger sequencing.

To delete the genome sequence around rs2280381, we utilized a dual-guide RNA strategy using two Cas9-guide RNA constructs. gRNAs were designed using an online tool (https://chopchop.cbu.uib.no/#), 1 pair of gRNAs around rs2280381 with the highest editing efficiency and a relatively lower off-target rate was chosen. gRNA oligoes were annealed and subcloned into the BbsI linearized px458 vector (Addgene,48138). 2×10^6^ cells were transfected with 5 μg px458-gRNA1 and 5 μg px458-gRNA2 plasmids using Neon system. Single cells with strong GFP signal were sorted into a 96-well plate by FACS. After 14 days of culture, genomic deletions were screened with Sanger sequencing of PCR amplicons. Electroporation conditions for each cell line were as follows: U-937, 1400 v, 10 ms, 3 pulses; Raji, 1350 v, 30 ms, 1 pulse; Jurkat, 1350 v, 10 ms, 3 pulses.

### CRISPRa screening

To design CRISPRa gRNAs, we first downloaded candidate SNP-centered 200 bp length sequences from human genome build GRCh38/hg19 (https://genome.ucsc.edu/cgi-bin/hgGateway) and utilized the CHOPCHOP online gRNA design tool (https://chopchop.cbu.uib.no/#) to obtain gRNA according to higher efficiency and lower off-target rate. For each candidate SNP, 3 gRNAs were designed around the SNPs and synthesized by GenScript Inc. The gRNA was dissolved to 35 μM concentration and stored at −20 °C. Prior to delivering gRNA into cells, U-937 cells stably expressing dCas9-vp64-Blast (Addgene 61425) and MS2-P65-HSF1-Hygro (Addgene 61426) were established by transduction of corresponding lentivirus following selection with 10 μg/mL Blasticidin (Invivogene, ant-bl-5) and 300 μg/mL Hygromcin (Thermo Fisher, 10687010) for one week. For screening, 2×10^5^ U-937 cells were resuspended in Buffer R, and 0.5 μL of each gRNA targeting the corresponding SNP were added into the cells. Then, the gRNA-Cells-buffer R mixture was aspirated into the 10 μl Neon pipette tip, and transfected using the Neon transfection system with the condition 1400 V, 10 ms, 3 pulses. After transfection, the cells were immediately transferred into a 24-well plate containing pre-warmed 10% FBS+90% RPMI-1640 media. After 24 hours culture, cells were collected to extract RNA.

### CRISPR SAM assay in the U-937 cell line

sgRNAs targeting the rs2280381-containing region were synthesized, annealed and cloned into lenti-sgRNA(MS2)-zeo backbone plasmid (Addgene, 61427) using restriction enzyme BsmBI (NEB, R0580L). sgRNA lentivirus particles were produced and transduced into a U-937 cell line stably expressing dCas9-VP64 and MS2-P65-HSF1 fusion proteins. Cells were selected with 400 μg/mL Zeocin (R25001, Thermo Fisher) for 72 h.

### PBMC isolation

Healthy human donors were recruited and signed informed consent according to the internal review and ethics boards of Renji Hospital, Shanghai Jiao Tong University. PBMCs were isolated using Ficoll-Paque density gradient solution (density =1.077 g/ml; GE Healthcare). Peripheral blood was mixed in a 1:2 ratio with phosphate-buffered saline (PBS) containing 2% FBS and 2 mM EDTA. After density gradient centrifugation (400 × g, 35 min, no brakes), the PBMC layer was carefully removed and the cell pellets were washed twice with PBS for further study.

### Lentivirus production

3×10^5^ HEK-293T cells were seeded into a six-well plate and incubated at 37 °C and 5% CO_2_ for overnight. Then cells were transfected with 1mg of targeting plasmid, 250 ng of pMD2.G (Addgene, 12259), and 750 ng of psPAX2 (Addgene, 12260) using 3 μL of Lipofectamine 2000 (Thermo Fisher, 11668-019). The media was changed after transfection for 6 hours. After transfection for 72 hours, virus supernatant was collected and centrifuged at 4 °C for 10 min to remove the debris. The supernatant was aliquoted, and stored at −80 °C.

### Circular chromatin conformation capture assay (4C) sequencing

To perform 4C-seq experiments, 1×10^7^ cells were collected and cross-linked by 1% formaldehyde solution. Then cells were quenched by 125 mM glycine solution. Cell pellets were resuspended in 5 mL cold lysis buffer (50 mM Tris, 150 mM NaCl, 5 mM EDTA, 0.5% NP-40, 1% Triton X-100, 1× protease inhibitor) and incubated on ice for 10 min. After lysis, cell nuclei pellets were collected, washed and resuspended in 500 μL 1× Csp6I buffer. 15 μL 10% SDS was added and incubated for 1 hour in a shaker at 750 r.p.m, then 75 μl 20% Triton X-100 was added and incubated for another 1 h with gentle shaking to sequester the SDS. 200 units of Csp6I enzyme (ThermoFisher, FD0214) (for rs2280381 view point, Csp6I was replaced with MboI (NEB, R0147M)) were added for a 4 h incubation at 37 °C in a shaker at 900 r.p.m. Then, 200 units of Csp6I enzyme was re-added and incubated at 37 °C in a shaker at 900 r.p.m overnight. Enzyme was inactivated at 65 °C for 20 min, 700 μL 10×T4 DNA ligase buffer was added and supplemented with Milli-Q ddH_2_O to a total volume of 7 mL. Then, 100 Units of T4 DNA ligase was added and incubated at room temperature for 6 hours. After that, 30 μL of Proteinase K (10mg/ml) was added and incubated at 65 ° for overnight. The remaining RNA was cleared by adding 30 μL RNase A (10mg/ml) and incubating at 37 °C for 45min. DNA was extracted with equivalent phenol/chloroform/isoamyl, and the pellets were dissolved in 150 μL 10mM Tris-HCl (pH 7.5). 50 μL of 10 ×NlaIII buffer and 50 units of NlaIII enzyme were added and supplemented with Milli-Q ddH_2_O to 500 μL volume. Samples were incubated at 37 °C for overnight. Enzyme was inactivated at 65 °C for 20 min, add 1.4 mL 10 × T4 DNA ligation buffer,100 Units of T4 DNA ligase, supplement Milli-Q ddH_2_O to 14 mL and ligate at room temperature for 4 h. DNA was purified using phenol–chloroform and further purified with the QIAquick PCR purification kit (Qiagen, 28106). The DNA concentration was detected by Qubit (ThermoFisher). The 4C-seq library was constructed by amplification of template using the 2×High-Fidelity Master Mix kit (Tsingke, TP001) with locus-specific primers containing Illumina sequences. The libraries were purified and sequenced on a HiSeq × ten (Illumina). 4C-seq data were analyzed using the software pipeline 4Cseqpipe (version 0.7), with settings: -stat_type median, trend_resolution 2000. Normalized trend was computed within the genomic region (chr16: 85,860,001– 86,060,000) for both viewpoints. Bowtiealign (version 1.2) was used to map captured reads to the Homo sapiens genome assembly GRCh37 (hg19) with the settings: -m 1 and captured fragments on chromosome 5 (reads per million more than 20) were listed.

### Cas9 RNP assembly

Alt-R crRNAs and Alt-tracrRNA-ATTO550 (IDT, 1075928) were ordered from Integrated DNA Technologies (IDT) and dissolved with Nuclease-Free Duplex Buffer (IDT) to 200 μM concentration. Equimolar concentrations of two oligos were mixed to a final 44 μM concentration and annealed. For each reaction, 22 pmol of crRNA-tracrRNA duplex and 18 pmol of HiFi Cas9 protein (1081061, IDT) were mixed in Buffer T to a final volume 1μL and incubated at room temperature for 10 min to prepare the Cas9 RNP.

### Primary immune cell subset isolation and editing

CD3+ T cells, CD14+ monocytes and CD19+ B cells were isolated from human PBMCs using the Human CD3+ T Cell Isolation kit (Miltenyi Biotec,130-050-101), Human CD14+ monocytes Isolation kit (Miltenyi Biotec, 130-050-201) and Human CD19+ B Cell Isolation kit (Miltenyi Biotec,130-050-301) respectively.

For T cell editing, after isolation, T cells were cultured in OpTmizer™ CTS™ T-Cell Expansion SFM medium (Thermo Fisher, A10458-03) supplemented with CD3/CD28 dynabeads (Thermo Fisher, 11131D) for 48 hours. Before transfection, CD3/CD28 dynabeads were removed and T cells were cultured for another 6 hours. Then 2 x 105 cells were washed twice with PBS and resuspended into 9 μL of Buffer T, mixed with Cas9 RNP and electroporated using the Neon transfection system with the condition 1400 V, 10 ms, 3 pulses. After that, T cells were transferred to the culture medium supplemented with 30 IU/mL IL-2 (Peprotech, 200-02A). After electroporation for 3 days, cells were collected to extract RNA and DNA.

For B cell editing, after isolation, B cells were cultured in RPMI-1640 medium with 10% (vol/vol) HI-FBS, 2 mM L-Glutamine, 55 μM β-mercaptoethanol, 50 IU/mL interleukin 4 (Peprotech, 200-04) and supplemented with CD40 ligand (Miltenyi Biotec, 130-098-775) for 48 hours. 1.2 x 10^5^ cells were collected and washed twice with PBS and resuspended into 9 μL of Buffer T, mixed with Cas9 RNP and electroporated using the Neon transfection system with the condition 1400 V, 10 ms, 3 pulses. After transfection, cells were immediately transferred to 500 μL of culture medium and cultured for 3 days, then cells were collected to extract RNA and DNA.

For monocyte editing, 2.5 x 10^5^ monocytes were washed twice with PBS and resuspended into 9 μl of Buffer T, mixed with Cas9 RNP, 1 μL Alt-R Cas9 Electroporation Enhancer and electroporated using the Neon transfection system with the condition 1600 V, 10 ms, 3 pulses. Cells were immediately transferred to 200 μL of medium containing 90% RPMI-1640 medium, 10% (vol/vol) HI-FBS, 2 mM L-Glutamine and 55 μM β-mercaptoethanol. After electroporation for 24 hours, cells with strong ATTO550 signal were collected to extract RNA and DNA.

### DNA methylation analysis

DNA was extracted from cells using the QIAamp DNA Mini Kit (Tiangen, DP304) and quantified using NanoDrop. Then DNA was treated with sodium bisulfite by using the EZ DNA Methylation-Gold Kit (Zymo, D5006) according to the manufacturer’s recommendations. Promoter methylation of *IRF8* was determined using quantitative methylation-specific polymerase chain reaction.

### DNA affinity precipitation assay (DAPA)-mass spectrometry

Cells were lysed to extract the nuclear lysates, and 200 μg nuclear extracts were mixed with 50 pmol of 5′-biotinylated DNA probes in the Buffer (20 mM HEPES, pH 7.9, 10% glycerol, 50 mM KCl, 0.2mM EDTA, 1.5 mM MgCl_2_,100 μg/mL Sheared Salmon sperm DNA, 1 mM dithiothreitol, and 0.25% Triton X-100) and incubated on ice for 45 min. Then Dynabeads™ M-280 Streptavidin (11205D, Thermo Fisher) was added and rotated for 2 hours at 4 °C. The enriched proteins were dissociated by the addition of 2 x Laemmli sample buffer (161-0737, Bio-Rad) and boiled at 95 °C for 10 min. The boiled protein samples were digested by trypsin for MS analysis.

### Liquid chromatography-MS/MS analysis and data processing

The tryptic peptides were dissolved in 0.1% formic acid (solvent A), and directly loaded onto a home-made reversed-phase analytical column (15 cm length, 75 μm i.d.). The gradient comprised an increase from 6% to 23% solvent B (0.1% formic acid in 98% acetonitrile) over 16 min, 23% to 35% in 8 min and climbing to 80% in 3 min then holding at 80% for the last 3 min, all at a constant flow rate of 400 nl/min on an EASY-nLC 1000 UPLC system.

The peptides were subjected to NSI source followed by tandem MS (MS/MS) in Q ExactiveTM Plus (Thermo) coupled online to the UPLC. The electrospray voltage applied was 2.0 kV. The m/z scan range was 350–1800 for full scan and intact peptides were detected in the Orbitrap at a resolution of 70,000. Peptides were then selected for MS/MS using NCE setting as 28 and the fragments were detected in the Orbitrap at a resolution of 17,500. A data-dependent procedure was used that alternated between one MS scan followed by 20 MS/MS scans with 15.0 s dynamic exclusion. Automatic gain control was set at 5E4.

The resulting MS/MS data were processed using Mascot Daemon (version2.3.0). Tandem mass spectra were searched against 2019-uniprot-human database. Trypsin/P was specified as cleavage enzyme allowing up to two missing cleavages. Mass error was set to 10 p.p.m. for precursor ions and 0.02 Da for fragment ions. Carbamidomethyl on Cys were specified as fixed modification and oxidation on Met was specified as variable modification. Peptide confidence was set at high and peptide ion score was set >20.

All oligoes used in this paper are listed in Supplementary Tables in Supplementary Information.

### Statistical analysis

All statistical analyses were performed using R Studio (version 1.0.136) with R version 3.3.3 and GraphPad Prism 7 software. Data are shown as mean ± SEM. ‘‘n’’ represents the number of technical replicates of the representative biological replicate unless otherwise mentioned. Details of the statistical analysis for each experiment can be found in the relevant figure legends. All statistical analyses were calculated using a paired or unpaired two-tailed Student’s t test as indicated in the figure legend unless otherwise mentioned. Asterisks define the significance level (* *P* ≤ 0.05; ** *P* ≤ 0.01; *** *P* ≤ 0.001).

## Data availability

The RNA sequencing data and 4C sequencing data that support the findings of this study have been deposited in the ArrayExpress database under accession codes <u>E-MTAB-10126</u> and <u>E-MTAB-10120</u> respectively. The public ATAC sequencing data <u>E-MTAB-8982</u> was used to analyze the chromatin accessibility. All other remaining data are available within the Article and Supplementary Files or are available from the authors upon request.

## Acknowledgment

This study was supported by grants from the National Natural Science Foundation of China (31930037, 31630021 and 81102266), National Human Genetic Resources Sharing Service Platform (2005DKA21300), Shanghai municipal key medical center construction project (2017ZZ01024-002), Shenzhen Science and Technology Project (JCYJ20180504170414637 and JCYJ20180302145033769), Shenzhen Futian Public Welfare Scientific Research Project (FTWS2018005) and Sanming Project of Medicine in Shenzhen (SZSM201602087).

## Author Contributions

N.S, GJ.H and T.Z designed the project. T.Z, GJ.H, XY.Z, YT.Z, YW.S and N.X performed the experiments, YT.Q performed the ATAC-seq experiment. XY.Z and C.Y performed analyzed the bioinformatics data. M.Z and Y.G analyzed the 4C-seq data. YF.W, WL.Y, YJ.T and T.Z analyzed the genetic data. J.B.H, B.N., K.M.K., L.C.K. and
M.T.W analyzed the genetic association data and revised the manuscript. HH.D, XL.Z, H.X and JY.M collected human samples and performed analysis. N.S, GJ.H and T.Z prepared the manuscript.

