## supplemental figure for "CRISPRa screen on a genetic risk locus shared by multiple autoimmune diseases identifies a dysfunctional enhancer that affects *IRF8* expression through cooperative lncRNA and DNA methylation machinery"

### Supplementary figures

**A**

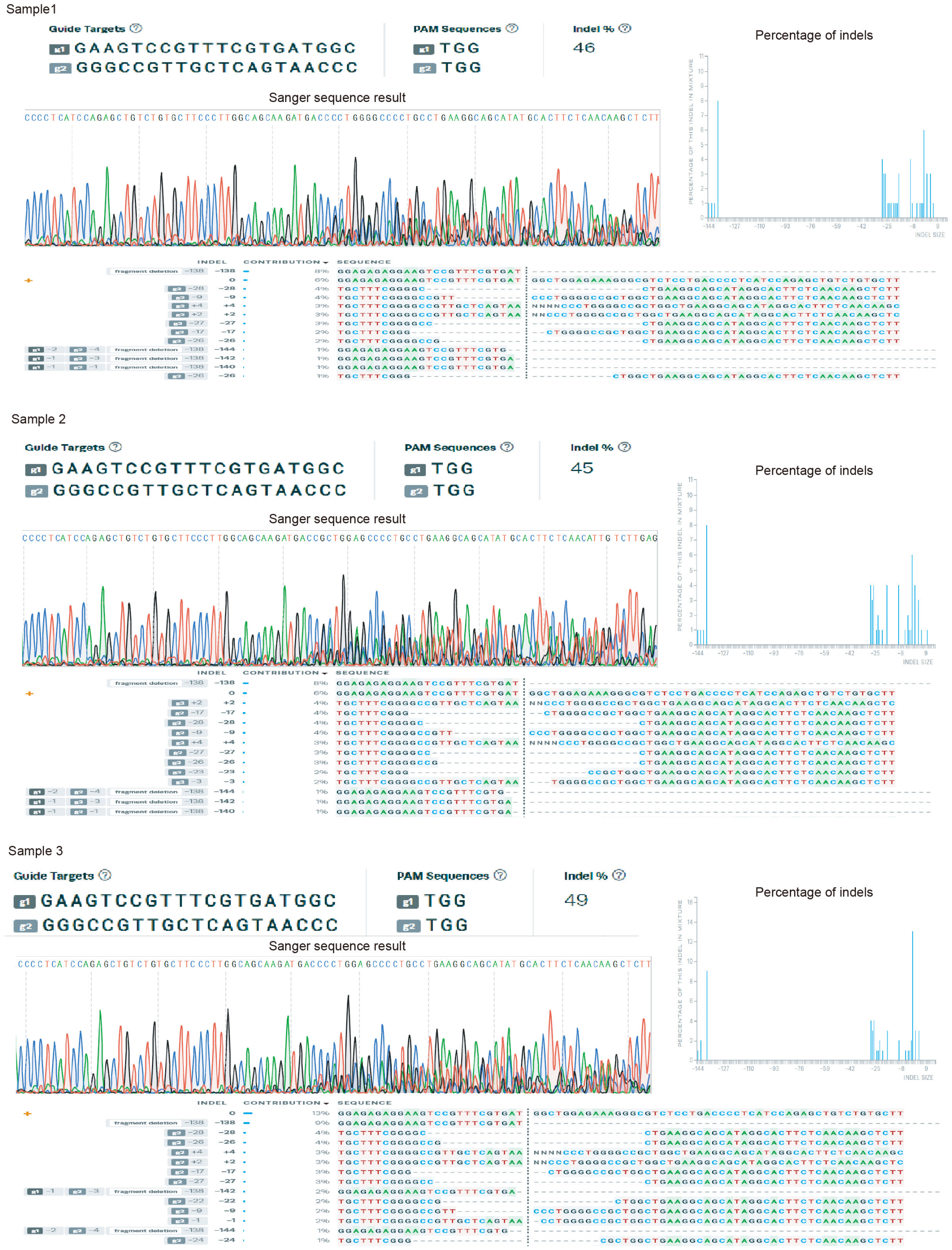

**B**

Sample1

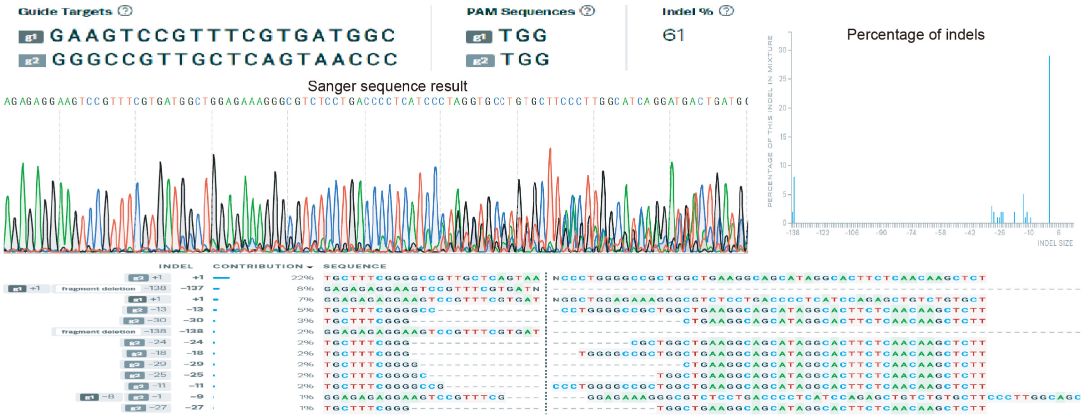


Sample2

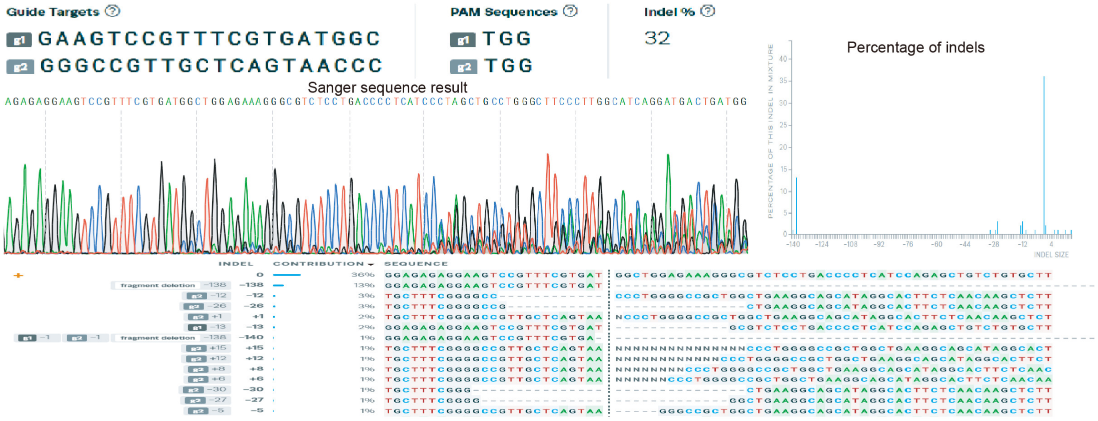


Sample3

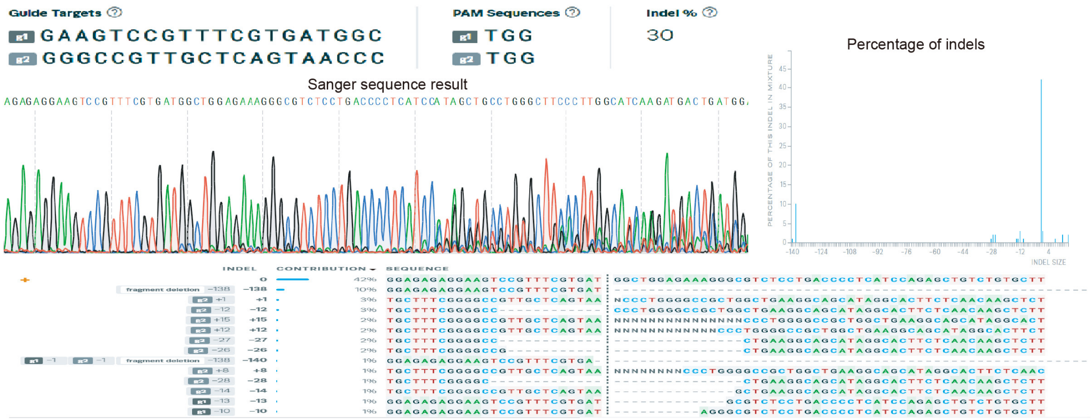

C

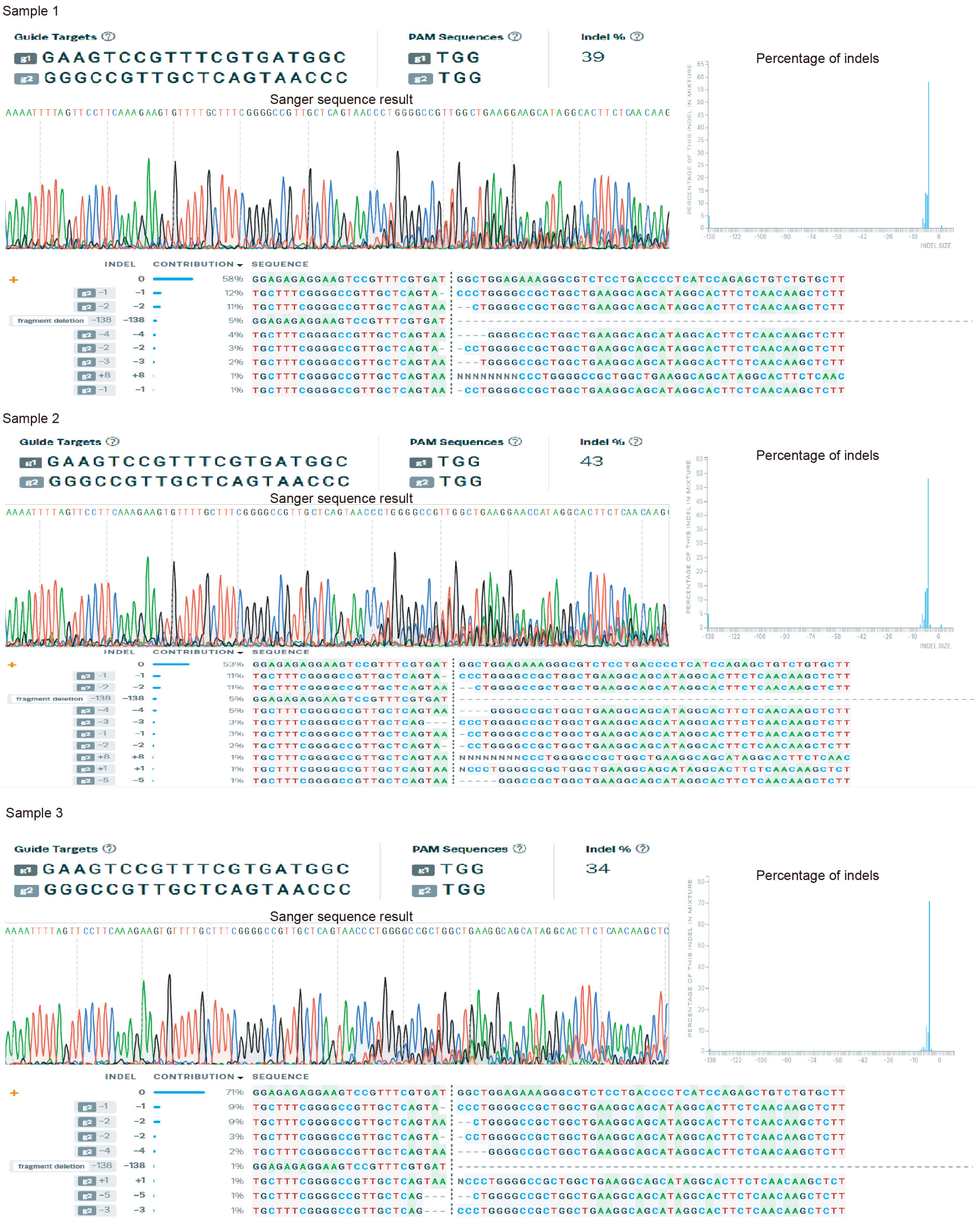


**Fig. S1 Related to Fig. 2.** Indel frequency in the edited immune cell subsets in vitro by ICE analysis. **(A)** Human CD3+ T cells. **(B)** Human CD19+ B cells. **(C)** Human CD14+ monocytes.

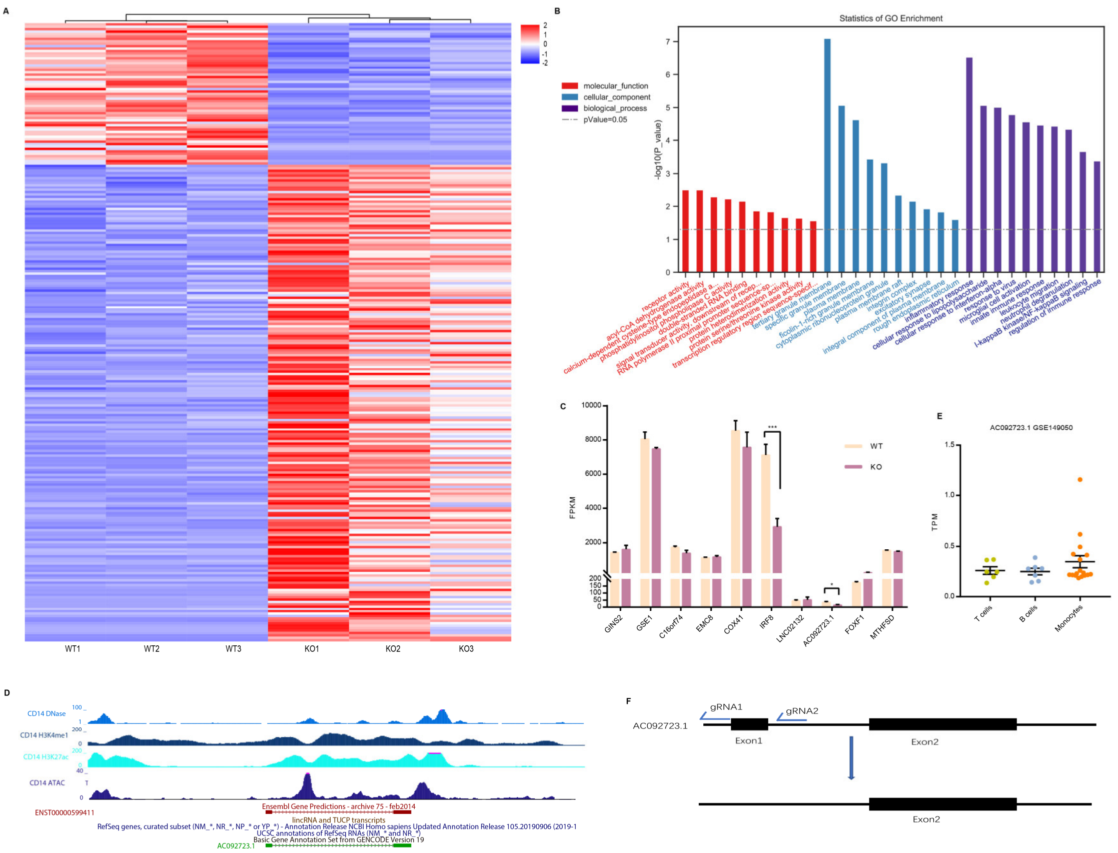


**Fig. S2 Related to Fig. 3. (A)** Heatmap of differentially expressed genes regulated by the rs2280381-containing region as measured by RNA-seq (log2 fold change >1.2 and *P* value <0.05). **(B)** Top 10 GO categories associated with differentially expressed genes. Categories were ranked by -log10 (*P* value). **(C)** RNA-seq analysis of the expression of genes within a 1-Mb window of rs2280381 in U-937 WT clones and KO clones (n = 3, biological samples replicates). **(D)** *AC092723.1* is located in an enhancer region with strong H3K4me1, H3K27ac, DNase and ATAC signals. **(E)** *AC092723.1* expression in different human immune cell subsets as measured by RNA sequencing from public database. **(F)** Deletion of the exon1 fragment of *AC092723.1* using CRISPR-Cas9.

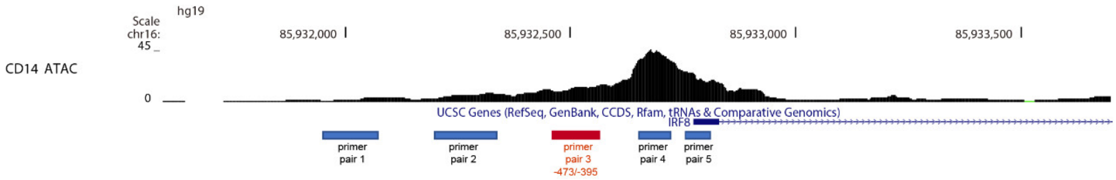


**Fig. S3 Related to Fig. 4.** The position of ChIRP primers in the *IRF8* promoter region.

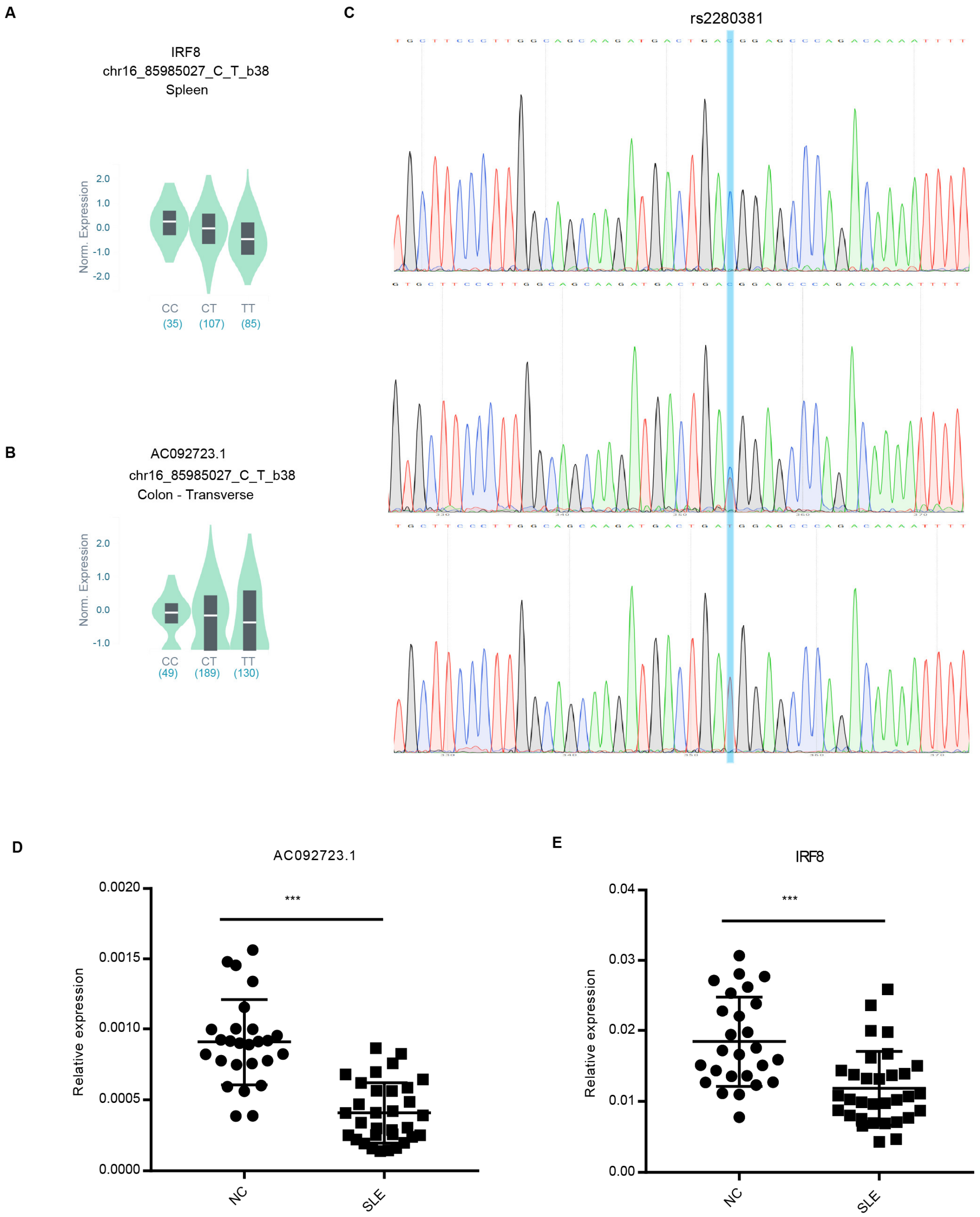


**Fig. S4 Related to Fig. 5. (A-B)** GTEx data analysis of the eQTL between *IRF8* or *AC092723.1* and rs2280381. **(C)** Genotype of clones was identified by Sanger Sequence. **(D-E)** RT-qPCR analysis of *AC092723.1* and *IRF8* expression in SLE patients. *P*-values are calculated using an unpaired two tailed Student's t-test. $*P < 0.05$; $**P < 0.01$; $***P < 0.001$.
